# A fluid dynamics-model system for advancing Tissue Engineering and Cancer Research studies: Dynamic Culture with the innovative BioAxFlow Bioreactor

**DOI:** 10.1101/2025.01.24.634745

**Authors:** Giulia Gramigna, Federica Liguori, Ludovica Filippini, Maurizio Mastantuono, Michele Pistillo, Margherita Scamarcio, Antonella Lisi, Giuseppe Falvo D’Urso Labate, Mario Ledda

**Affiliations:** Consiglio Nazionale delle Ricerche, Istituto di Farmacologia Traslazionale, Roma, via del Fosso del Cavaliere100; Italy; Cellex S.r.L, Roma, Piazzale delle Belle Arti, 2; Italy

## Abstract

In this study, we test an innovative bioreactor, particularly suitable for tissue engineering applications, named BioAxFlow. Unlike traditional bioreactors, it does not rely on mechanical components to agitate the culture medium, but on the fluid-dynamics generated thanks to the unique geometry of the culture chamber. The flow generated within ensures continuous medium movement, promoting consistent cell exposure to nutrients and growth factors. Using the human osteosarcoma cell line SAOS-2, the bioreactor’s ability to enhance cell adhesion and proliferation on polylactic acid scaffolds, mimicking bone tissue matrix architecture, is tested. The findings show that the bioreactor significantly improved cell adhesion and growth compared to static cultures, promoting a homogeneous cell distribution across the scaffold surfaces, which is crucial for developing functional tissue constructs. The bioreactor preserves the osteogenic potential of SAOS-2 cells as assessed by the expression of key osteogenic markers. Additionally, it retains the tumorigenic characteristics of SAOS-2 cells, including the expression of pro-angiogenic factors and apoptosis-related genes. These results indicate that the BioAxFlow bioreactor is an effective platform for tissue engineering and cancer research, offering a promising tool for both regenerative medicine applications and drug testing.

## 1 Introduction

The human bone is a highly dynamic tissue, which undergoes continuous structural changes: the so-called remodelling. Remodelling entails continuous interplay between all the bone cells, in diverse differentiation stages, such as resident mesenchymal stem cells (MSCs)-derived osteoprogenitors, osteoblasts, osteoclasts, osteocytes, monocyte/macrophage cells and T cells (1). An imbalance among bone cellular populations’ activity is representative of both aging and pathology. Indeed, aging is defined by several biological hallmarks, including resident stem cells depletion, tissue inflammation, ECM alterations, cellular senescence and metabolic dysfunction (2), while osteoporosis, a disease whose incidence increases in older people, is characterized by low bone mass and architectural deterioration, leading to enhanced bone fragility and increased fracture risk.

The operational definition of osteoporosis is based on bone mineral density (BMD) measurement (3), reflecting the porosity of the bone structure. For instance, in cortical bones, porosity is around 4% in young healthy adults and increases up to around 12% in people aged 60(4), whereas it can get to ca. 50% in people older than 60 (5). Nevertheless, osteoporotic fractures mostly occur in predominantly trabecular bones (i.e. proximal femurs, spines and distal radii), where they exert a critically important role in load transmission and energy absorption (6, 7). All the research conducted in the osteoporosis-field over the last decades strongly shows a correlation between BMD and the strength of the trabecular bone, as reflected by the elastic modulus calculated from micro computed tomography images (µCT) (8).

Bone fragility associated with osteoporosis represents a socio-economic burden on societies world-wide, as it is the root cause of most bone fractures. As an example, in France, Germany, Italy, Spain, Sweden and the UK, 2.68 million bone fractures are registered yearly. Such a figure is expected to increase 23% by 2030 (https://www.iofbonehealth.org/). Only in these countries, in 2017 fracture-related costs summed up to €37.5 billion.

These numbers suggest that novel and effective therapeutic approaches should be developed to break up the cost spiral and improve patients’ quality of life. Currently, animal models are used to study bone biology and mechanobiology or to test drugs for bone diseases. They are costly, complex, and often fail to capture human-specific features and to predict the clinical outcome (9). Their use is ethically questionable and will soon be reduced to comply with the 3R’s EU directory: Replacement, Reduction, Refinement. Many alternative cell-based in vitro models have been proposed, e.g. a tissue engineering (TE) approach would be desirable to the preparation of both sustainable and reliable bone substitutes and in vitro bone models.

Therefore, we aimed to develop a 3D platform mimicking the human bone tissue in vitro, able to recreate a microenvironment as close as possible to the native tissue, through means of the so-called TE-triad (10): 1) signals, 2) biomaterials and 3) cells. For ensuring that the signals, either chemical or mechanical, would have been homogenously distributed among the cultured cells, we have developed an innovative bioreactor, named BioAxflow (BAF), able to gently mix the cell culture medium without the use of mechanical parts (such as impellers or rotating walls) which, in turn, induce a high shear stress on the cultured cells.

In a tentative to achieve a gentle and homogeneous mass transfer within a culture chamber, microgravity-based technologies have been explored in the past. These technologies have been patented for application to bioreactor devices (i.e. US5846817A) in the 90’s but the use of rotating mechanical components introduce several limitations to the system. It includes a vertical chamber in which a set of rotating filters arranged in a central position interact with flexible membranes. In this bioreactor, the agitation and mixing process is based on fluid dynamics and therefore depends largely on the speed of rotation. The bioreactor we used (pending patent application), on the other hand, can gently mix the culture medium thanks to the fluid dynamics induced by the chamber-architecture, combined with the use of a peristaltic pump. Therefore, this technology is not dependent neither on microgravity nor mechanical parts, which enables the bioreactor we used to be an easy-to-use device and a breakthrough innovation in the 3D-dynamic cell culture field. Historically, bone-like inorganic biomaterials (e.g., hydroxyapatite or ß-tricalcium phosphate (ßTCP)), natural (e.g., ECM-based collagen type I, proteins) and synthetic polymers (e.g., polylactic-co-glycolic acid, poly (caprolactone (PCL)-co-L-lactide) have been used to mimic the ECM of healthy bone (11), hence to make the second element of the TE triad: the biomimetic scaffolds. Inorganics are osteoconductive and have similar stiffness to bone but are brittle. Natural polymers suffer from high batch-to-batch variability and may elicit immune reactions. Most polymers lack mechanical strength to withstand mechanical loading. Composite materials (e.g., fibre/filler-reinforced polymeric matrices) have been recently proposed to overcome such limitations (12). 3D scaffolds are generally prepared as mm-sized porous cylindrical plugs, yet it is difficult to prepare them with controlled and reproducible micro to macroscale architecture and scaffold morphology and architecture from nano- to microscale affects cell behaviour. To test the BAF bioreactor in the context of bone-tissue engineering, human osteosarcoma SAOS-2 cells were chosen as an osteoblast-like cellular model and used as the third player of the tissue engineering (TE) triad. The SAOS-2 cell line is recognized as one of the most representative osteoblast-like cell models for studying cell-material interactions in TE (13, 14). These cells exhibit an osteoblast phenotype with features closely resembling those of human primary osteoblasts (15). Compared to other osteoblast-like cell lines, SAOS-2 cells better mimic primary human osteoblast behaviour when interacting with biomaterials (16). These cells are characterized by high levels of Alkaline Phosphatase (ALP) activity, a hallmark of osteoblast differentiation, and exhibit growth factor expression and collagen production comparable to human primary osteoblasts (17). They express bone-specific markers, such as Collagen Type I, ALP, Osteocalcin (OCL), and Osteopontin (OPN), and produce bone matrix proteins. This makes them a preferred model for studying osteogenic differentiation and bone tissue dynamics. Additionally, SAOS-2 cells are widely used in osteosarcoma and bone-related research due to their osteoblastic properties and ability to replicate key aspects of bone biology and cancer pathology. SAOS-2 cells exhibit rapid proliferation and are resistant to certain apoptotic stimuli, making them ideal for studying the growth and survival of osteosarcoma cells under various conditions, including the effects of potential therapeutic agents. They also exhibit features of tumour progression, such as angiogenesis and metastatic potential and secrete Vascular Endothelial Growth Factor (VEGF), which supports new blood vessel formation, a critical factor in tumour growth and metastasis (18). Studies have highlighted the potential of targeting VEGF signalling in osteosarcoma, as its inhibition not only reduces angiogenesis but also promotes apoptosis, suggesting a promising therapeutic approach. This is especially relevant since VEGF overexpression is linked to worse prognosis in osteosarcoma patients (19). This cell line is frequently employed to investigate the effects of chemotherapeutic agents or targeted therapies on osteosarcoma, focusing on pathways regulating apoptosis (e.g., BAX/BCL2) and angiogenesis (e.g., VEGF inhibition)(20, 21). They provide a valuable platform for assessing drug resistance, sensitivity, and the molecular mechanisms underlying these processes.

In this study, we propose a novel strategy for 3D cell growth by culturing SAOS-2 cells within the BAF bioreactor to evaluate their adhesion and proliferation on bone-mimicking scaffolds. These experiments aimed at assessing the compatibility and effectiveness of the bioreactor system for supporting osteoblastic cell growth and scaffold colonization, a critical step in advancing bone tissue engineering strategies. The results demonstrate that the BioAxFlow bioreactor is a valuable platform for studying and developing functional tissue constructs in bone tissue engineering, as well as a useful tool for osteosarcoma research and cancer therapy testing.

## 2 Material and Methods

### 2.1 Bioreactor Setup

The bioreactor system (BioAxFlow by Cellex) comprises four main components: (1) a base (height 8 cm, Figure 1A) integrating inlet and outlet ports for controlled media flow, (2) a cylindrical chamber (diameter 8 cm, Figure 1A,B), (3) scaffold stand for scaffold placement (Figure 4 for an individual representation), and (4) a cap with two openings for vent caps and two sampling ports. All components were sterilized and assembled under the laminar flow hood and connected with sterile silicone tubing to a peristaltic pump (Watson Marlow 505 S) at the CNR-IFT laboratories (Figure 1B).

**Figure 1:**
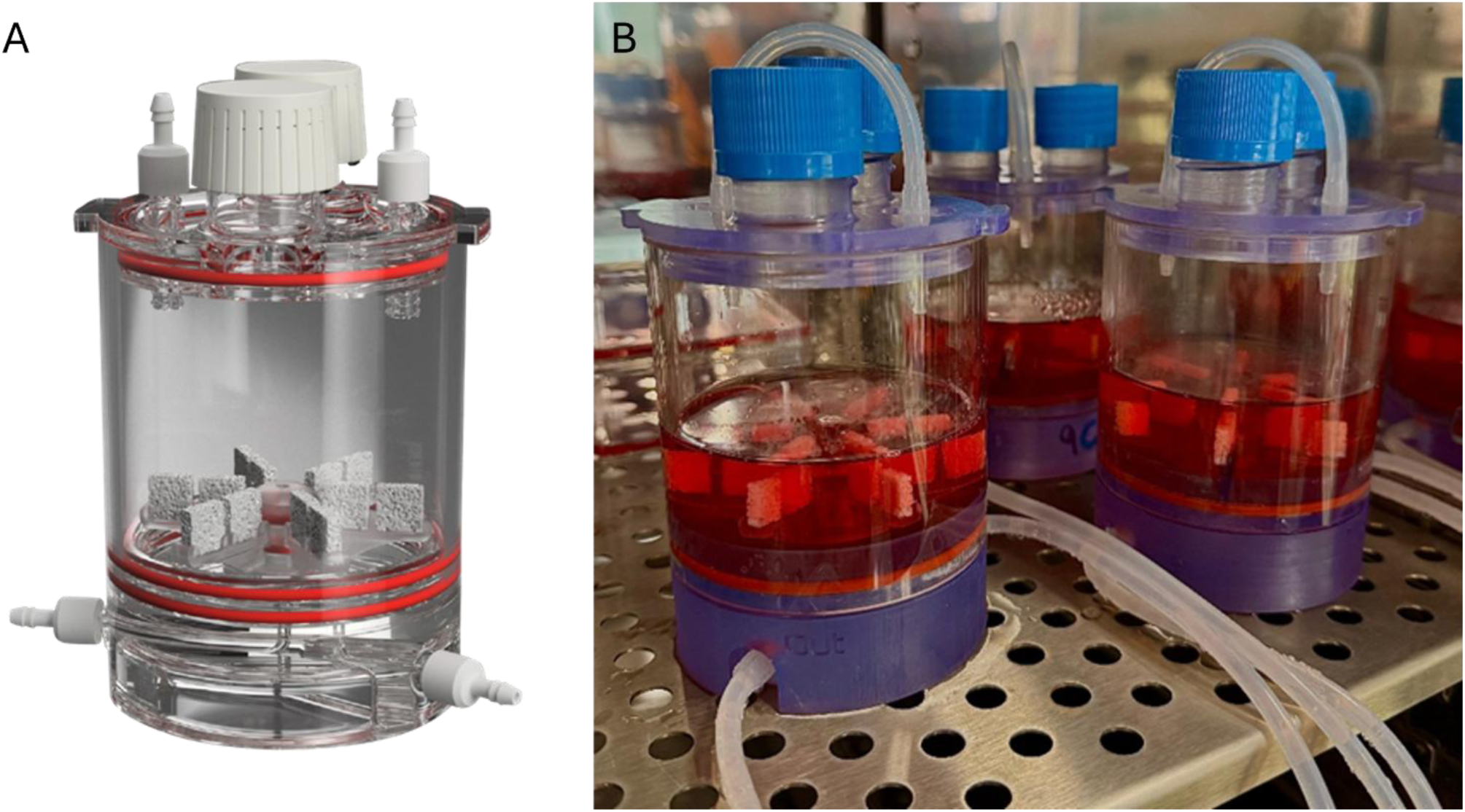
Rendering of BioAxFlow (A); BioAxFlow bioreactors to conduct two experiments in parallel, placed inside the CO_2_ incubator (B)

For the seeding process, a cell suspension was injected into the main chamber where scaffolds were positioned on the scaffold stand. The peristaltic pump was then activated to initiate cell seeding, an automated process designed to maximize cell use and ensure even cell distribution across scaffolds. The scaffolds are continuously surrounded and perfused by a recirculating cell culture medium in a closed-loop system (total volume: 110 mL). Media enter the bioreactor from an inlet port at the base and exit through an inner outlet, with both inlet and outlet ports featuring an internal diameter of 1.6 mm. The entire system is compatible with placement in a standard humidified incubator at 37°C and 5% CO₂.

### 2.2 Fluid dynamics simulations

The computational modelling carried out in COMSOL focusses on recapitulating the fluid dynamics characteristics of the medium flow, by investigating the velocity profiles, pressure, and oxygen distribution within the system under consideration. The model did not include the scaffolds and their holders, so that the computational efficiency was increased, and focussed on the macroscale interactions, hence allowing for the examination of mixing and oxygenation phenomena, both crucial for cellular proliferation and activity. The greatest influence on cellular proliferation and activity was played by the configurational and operational parameters set, rather than any scaffolds configuration, which had little role on the overall performance. Thus, the experimental data obtained from cells cultured on the scaffolds can add information to the parameters derived from the computational simulations and provide a validation of the expected conditions, as it has been shown later on. Although including both the scaffolds and their stand into the simulations would have enabled the estimation of more local effects, the approach used represents a reasonable trade-off between precision and complexity that fully meets the goals of the study.

More specifically, to model the media perfused within the BioAxFlow bioreactor, COMSOL Multiphysics® 6.3 was employed to streamline the construction of the initial simulation. The process began by defining global parameters, including the vessel’s diameter and height, as well as the internal channels and baffles in the base. Next, the fluid material was selected, with properties such as density and viscosity set to mimic water (1000 kg m^-^³ and 0.001 Pa s, respectively).

The key step involved specifying the physics of the model. This included defining the fluid domain and setting boundary conditions for both the chamber’s walls and the flow’s inlet and outlet. Additionally, initial values (such as for pressure or velocity) and free surface interactions were incorporated to accurately represent the system’s fluid dynamics.

The initial pressure and the initial velocity in all directions was set to zero. The motion of the fluid is governed by the Navier-Stokes equations, which can be seen as the application of Newton’s second law to fluid dynamics. For a compressible Newtonian fluid (Equation1), these equations take the form:

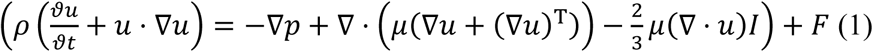

where *u* represents fluid velocity, *p* is fluid pressure, *ρ* is fluid density, and *μ* is the fluid dynamic viscosity. The terms in the equation correspond to inertial forces (first term), pressure forces (second term), viscous forces (third term), and external forces acting on the fluid (fourth term). The Navier-Stokes equations, developed by Navier, Saint-Venant, Poisson, and Stokes between 1827 and 1845, are always solved in conjunction with the continuity equation (Equation 2):

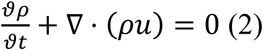

The Navier-Stokes equations represent the conservation of momentum, while the continuity equation ensures the conservation of mass. Together, they form the foundation of fluid flow modelling. By solving these equations with specific boundary conditions—such as inlets, outlets, and walls—it is possible to predict the fluid’s velocity and pressure within a given geometry (22). These solutions are crucial for understanding and optimizing fluid dynamics in systems like the BioAxFlow bioreactor.

The next step in constructing the model involved setting up the mesh (Figure 2), or finite element network, necessary to solve the defined boundary conditions. A physics-controlled mesh with normal element size was used to minimize computational demands, such as Random Access Memory (RAM) and simulation time. The finite element method converts the problem into a system of algebraic equations, yielding approximate values for the unknowns at discrete points within the domain. This approach divides the large problem into smaller, simpler parts called finite elements. The equations for each element are then assembled into a larger system that models the entire problem (23).

**Figure 2:**
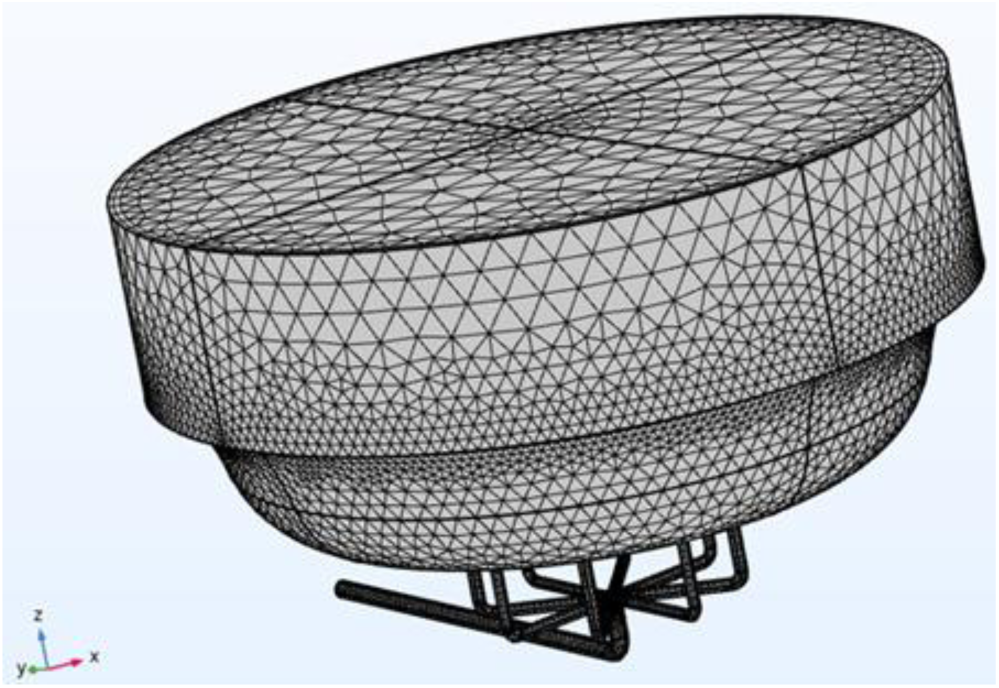
BioAxFlow (Volume = 110 mL) meshed geometry

The final step in the model setup involved selecting the appropriate study type, which also computed the initial conditions for time-dependent flow simulations within the vessel. Once all these steps were completed correctly, the computation could begin, with the process potentially taking several hours depending on available computational power.

### 2.3 Simulation of Oxygen distribution

The CFD models described previously were used to simulate the spatio-temporal distribution of oxygen. Used parameters are listed below:

D_O2_, O_2_ diffusion coefficient in aqueous media: 3×10^-^⁹ m² s^-1^ (24)
Cell-normalized O_2_ consumption rate in Saos-2 cells : 2 nmol min^-1^ 10^-^⁶ cells (25)
K_m_, Michaelis-Menten constant: 5.6 mmHg (26)
C_0_, O_2_ concentration at air-liquid interface: 0, 214 mol m^-3^ (24)
K_O2_, Henry’s law constant: 932.4 atm mol^-1^ L^-1^ (24)

Mass transport was by advection and diffusion (Equation 3), as described here:

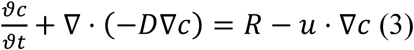

where c is the oxygen concentration, D is oxygen diffusion coefficient (m^2^ s^-1^), R is the reaction rate (mol m^-3^ s^-1^), and u is 3D velocity field (m s^-1^) (27). The concentration of oxygen in the media at t=0 was assumed to be in equilibrium with the concentration of oxygen in the air, as described by Henry’s law (Equation 4):

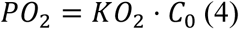

where KO_2_ is the Henry’s law constant and PO_2_ is the partial pressure of oxygen in the air (refer to the parameters listed above, including references) and C_0_ represents the concentration of oxygen in the media entering the bioreactor from the free surface. In the presence of cells, local oxygen concentrations would depend on mass transport of oxygen in the bioreactor and the rate of oxygen consumption by the cells. The rate of oxygen consumption by the cells was assumed to follow Michaelis–Menten (MM) kinetics (Equation 5):

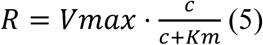

Where V_max_ in the maximal oxygen consumption rate (moles m^-3^ s^-1^), K_m_ in the MM constant (moles m^-3^). Refer to the parameters listed above, including references. A cell-normalized oxygen consumption rate (SAOS-2 cells) of 2 nmol min^-1^ 10^-^⁶ was used, along with a km value of 5.6 mmHg. The total number of cells in BioAxFlow (d=8 cm) was 11 million cells. V_max_ values were calculated by multiplying the cell-normalized oxygen consumption rate by the number of cells in the bioreactor and then dividing by the volume of the vessel (110 mL).

### 2.4 Scaffold Fabrication

Bone inorganic matrix mimicking scaffolds have been fabricated according to the procedures described in Zenobi et al., 2023 (28) using a percentage porosity of 52.5 % for reproducing the healthy bone condition. The scaffolds were designed using Meshmixer 3.5 software (v.2018, Autodesk, San Rafael). Briefly, the process began with a solid of 10 × 10 × 3 mm³. A three-dimensional random cluster of spheres was subtracted from this solid to create a porous structure. The sphere diameter (pore size) and centre-to-centre distance (spacing) values were set at 700 μm (28). The printing was conducted with a Prusa MK3S printer, and the chosen printing material was a commercially available polylactic acid (PLA) filament, which had a 1.75 mm diameter and was sourced from FILOALFA. This PLA filament was carefully heated to a precise extrusion temperature of 205°C, allowing it to flow through a nozzle with a diameter of 0.4 mm. During the printing process, the printer bed was consistently maintained at a temperature of 60°C to ensure optimal adhesion and stability. Importantly, it is worth noting that all samples underwent fabrication using identical printing parameters, thereby guaranteeing the homogeneity and standardization of the produced scaffolds and control specimens. The dimensions obtained for the parallelepiped scaffolds were 10 mm × 10 mm × 3 mm (Figure 3).

**Figure 3:**
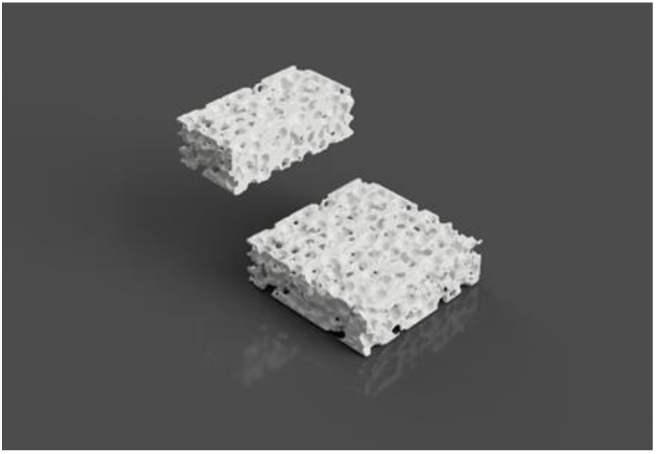
Rendering of PLA scaffold

### 2.5 Scaffold stand

During the scaffold stand fabrication procedure, a design approach aimed at optimizing the allocation of scaffolds within the bioreactor was adopted. The structure was conceived to simultaneously support twelve parallelepiped scaffolds (10×10×3 mm), ensuring efficient use of the available room. The scaffold stand, as well as the base and cap of the bioreactor, were designed using Fusion 360, a powerful CAD software that facilitates the creation of complex geometries and allows for precise modifications based on functional requirements Figure 4. Each scaffold is designed to maximize cellular interaction and nutrient flow, which is essential for the success of cell culture processes. Moreover, the configuration of the scaffold stand facilitates the insertion and removal of scaffolds from the bioreactor chamber, ensuring a homogeneous distribution of environmental parameters, such as temperature and oxygen.

**Figure 4:**
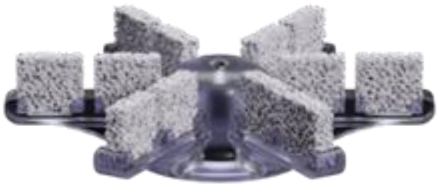
Rendering of the developed scaffold stand.

### 2.6 Cell Culture

The human osteosarcoma SAOS-2 cell line (ATCC, HTB-37, Rockville, MD, USA) was cultured in high-glucose Dulbecco’s modified Eagle’s Medium (DMEM; Euroclone), supplemented with 10% heat-inactivated fetal bovine serum (FBS; Euroclone), 2 mM L-glutamine (Sigma), 1.0 unit·ml^-1^ penicillin (Sigma), and 1.0 mg ml^-1^streptomycin (Sigma). Cells were maintained in plastic Petri dishes at 37°C in a humidified incubator with 5% CO₂. For experiments, polylactic acid (PLA) scaffolds were sterilized by immersion in 70% ethanol for 30 minutes, rinsed with phosphate-buffered saline (PBS), and placed within the BAF bioreactor, using an appropriately printed stand to accommodate the parallelepipedal scaffolds, as described above (for dynamic culture conditions) or placed in plastic multi-well plates (for static culture controls). SAOS-2 cells were cultured at a concentration of 150,000 cells ml^-1^ or 50,000 cells ml^-1^ directly inside the bioreactor chamber (where PLA scaffolds were placed into) or on PLA scaffolds in multi-well plates. Cultures were maintained for up to 10 days.

### 2.7 Cell adhesion and growth analysis

Cell adhesion and growth trend was quantified by Trypan Blue exclusion assay. SAOS-2 cells cultured within the bioreactor (dynamic conditions) or seeded onto PLA scaffolds in multi-well plates (static conditions) were harvested with 0.1% trypsin–EDTA (Sigma), washed twice with PBS and the total number of nucleated and viable cells was counted by Trypan Blue dye (0,4%) (Sigma) exclusion assay using a Bürker hemacytometer chamber. This protocol was performed at 1, 4, 7 and 10 days. Each experiment was repeated three times.

### 2.8 Real-Time quantitative RT-PCR analysis

Total RNA was extracted from SAOS-2 cells cultured on PLA scaffolds under dynamic (bioreactor) or static conditions. Cells were harvested with 0.1% trypsin–EDTA (Sigma), washed twice with PBS, and processed after 4, 7, and 10 days. RNA isolation was performed using TRIzol Reagent (Invitrogen) according to the manufacturer’s protocol. First-strand cDNA synthesis was performed with 1 µg of total RNA using random primers and the iScript™ cDNA Synthesis Kit (Bio-Rad). Gene expression was assessed by RT-qPCR using SsoAdvanced™ Universal SYBR® Green Supermix (Bio-Rad) on a Bio-Rad Real-Time PCR Detection System. Reactions were conducted in triplicate, with each 20 µl reaction containing 0.5 µl cDNA template and primers at concentration of 250 nM. The primers used are listed in Table 2.

Amplifications followed these cycling conditions:

50°C for 2 minutes,

95°C for 10 minutes (DNA polymerase activation),

40 cycles of 95°C for 15 seconds, and 60°C for 1 minute.

Melting curve analysis, performed using Bio-Rad Dissociation Curves software, confirmed product specificity. Negative controls, omitting RNA or reverse transcriptase during cDNA synthesis, were included to rule out contamination. Relative gene expression was normalized to GAPDH as an endogenous control, and data were analysed using the 2^−ΔΔCt method as described by Livak and Schmittgen (29). Amount of target was calculated using the 2-ΔΔCt equation.

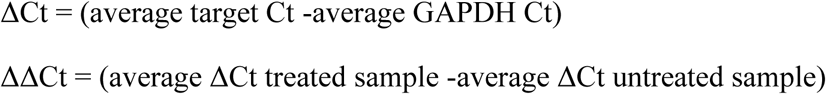

Before using ΔΔCt method for quantification, we performed a validation experiment to demonstrate that amplification efficiency for the target genes and the reference GAPDH gene were equal. The primers were designed using the GeneRunner software and purchased from Eurofins, their respective sequences used are reported in Table 1.

**Table 1.**
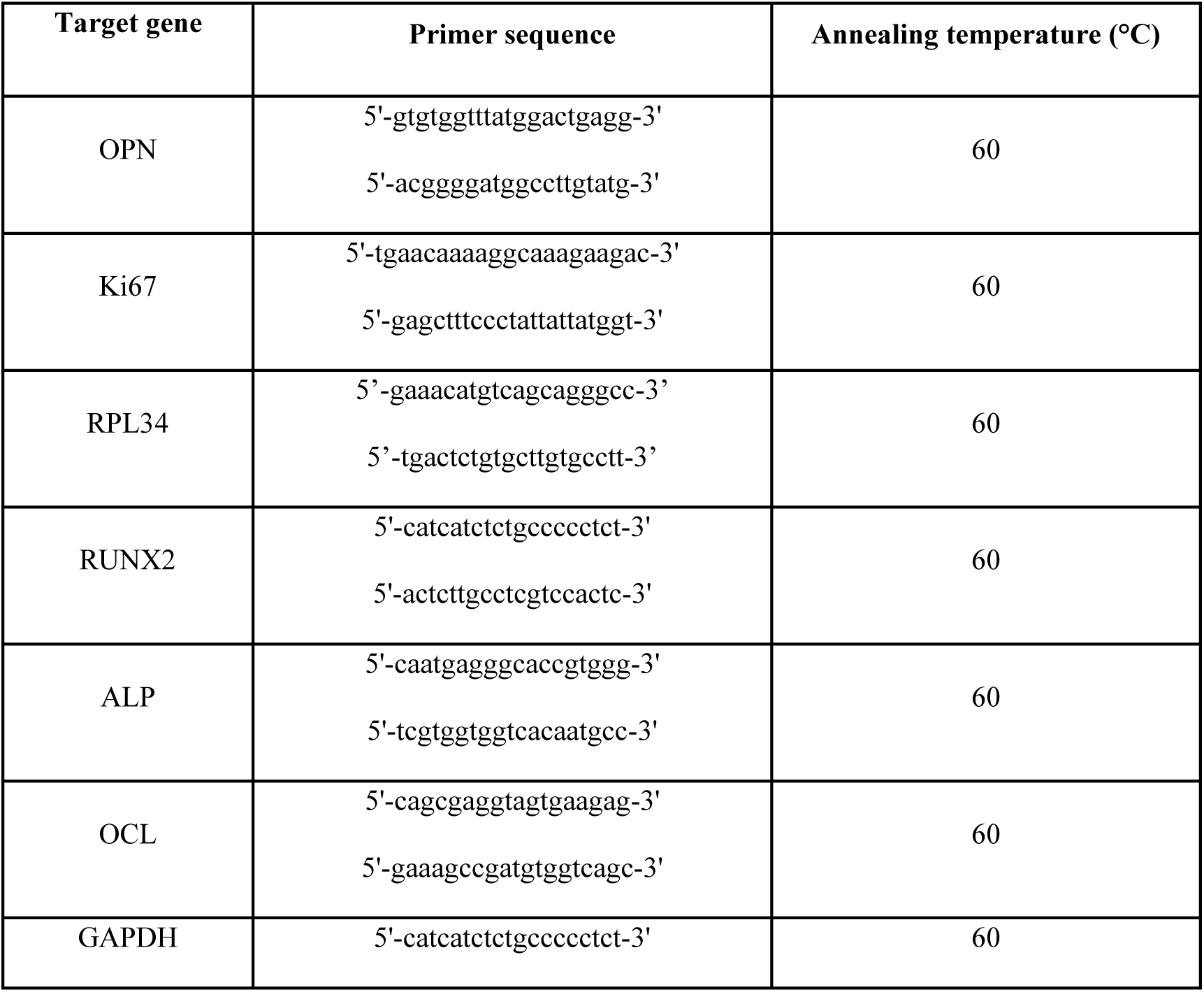
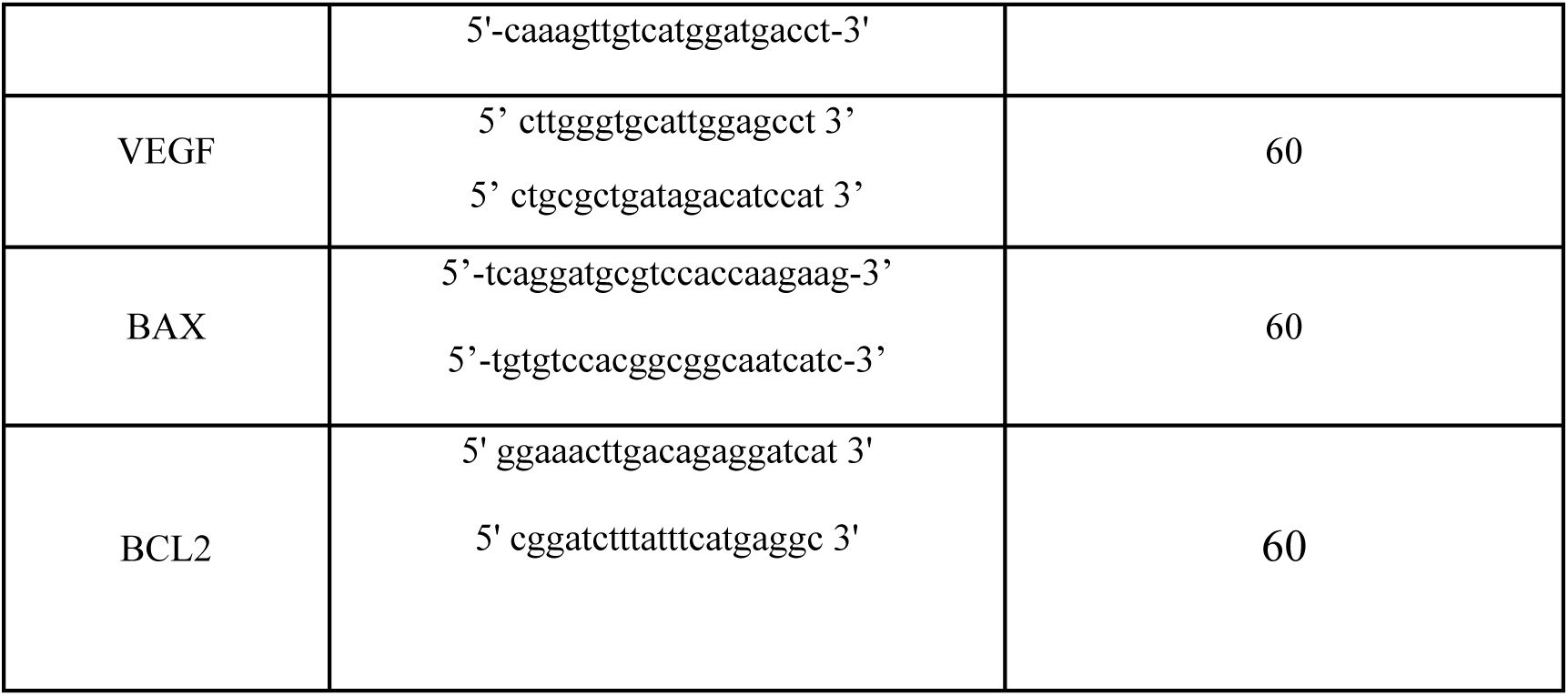
Sequence of primers used for qRT-PCR.

### 2.9 Confocal Laser Scanning Microscopy

SAOS-2 cells cultured for 24 h on PLA scaffolds under dynamic (bioreactor) or static conditions were repeatedly washed with PBS and fixed in 4% paraformaldehyde for 15 min and rinsed twice with PBS. The cells were stained for nuclei localization with propidium iodide (2 μg ml^-1^; Sigma). The fluorescence analyses were performed by using LEICA TCS 4D Confocal Microscope. Confocal optical Z sections were acquired at 5-μm intervals for entire surface of the scaffolds and a middle confocal Z section is shown.

### 2.10 Statistical analysis

MedCalc software was used for statistical analysis, and the significance level adopted for all analyses was p < 0.01. For the results of cell adhesion assay, data were analysed by Student’s T-test. For cell growth analysis data were analysed by two-way ANOVA-test, in which one factor is the type of treatment (Static, BAF, Static to BAF and BAF to Static) and the other factor is time (1 day, 4 days, 7 days and 10 days).

## 3 Results

### 3.1 BioAxFlow fluid Dynamics Simulations

The fluid dynamics simulations were performed using a flow rate of 80 mL min^-1^, and the results highlight key aspects of fluid behaviour within the bioreactor under perfusion conditions.

#### 3.1.1 Velocity Field

The visualization of the velocity field model characterization provides an insight into how the geometry of the bioreactor influences fluid mixing dynamics. A cut-plane along the ZX-plane was used to create a cross-sectional view of the reactor, allowing for detailed observation of internal flow patterns. For clarity, all walls of the model were rendered invisible. The combined streamline plot revealed the directionality and intensity of fluid motion throughout the volume, indicating a high degree of mixing efficiency. Notably, the velocity of the fluid was higher in the inlet and outlet flow channels, as well as at the bottom of the vessel. This behaviour is consistent with the design of the bioreactor, which promotes thorough mixing to ensure homogeneity in nutrient distribution and waste removal during perfusion (Figure 5A).

**Figure 5:**
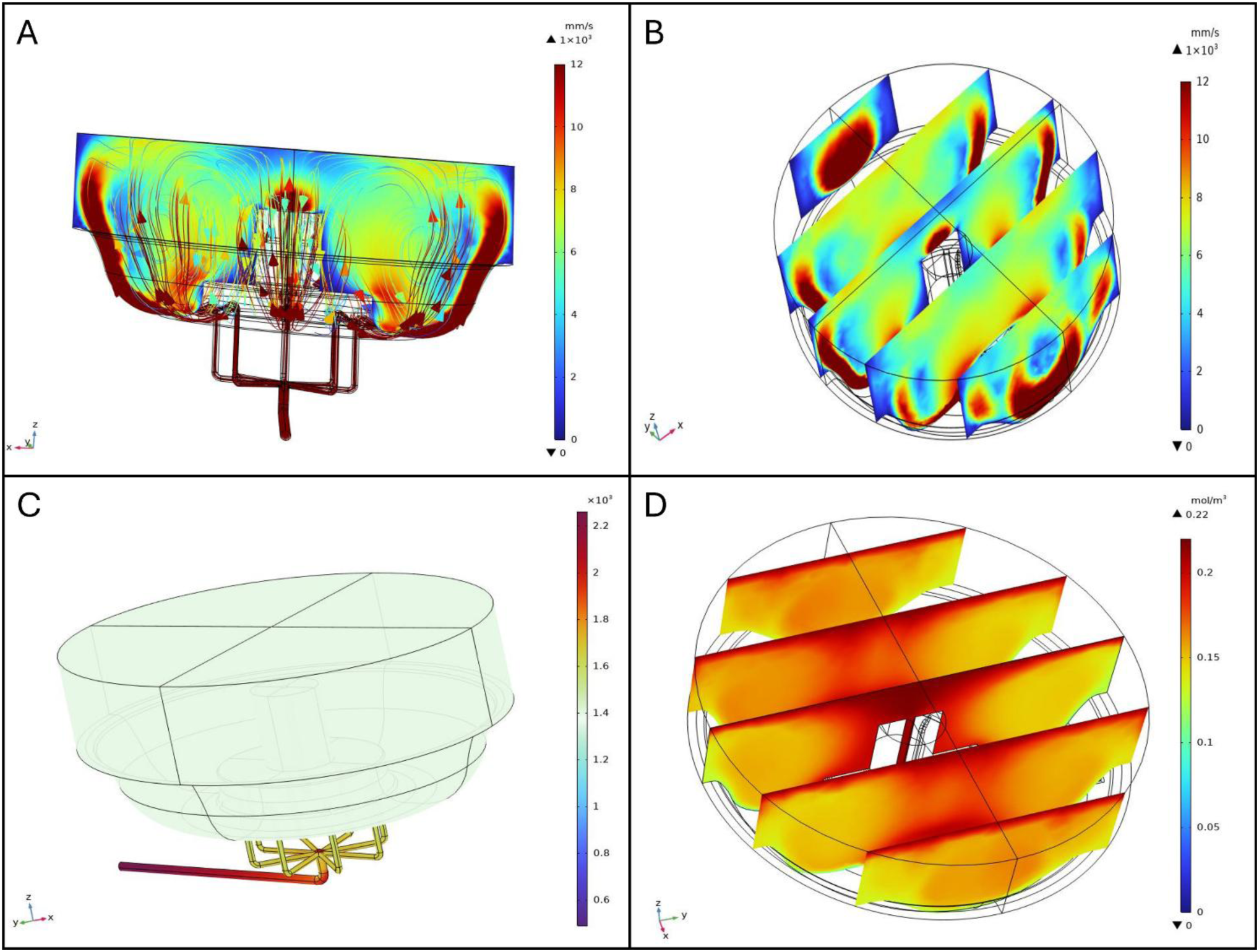
Fluid dynamics simulations: (A)and streamline profiles, (B) Slice plot in ZX-plane (distributed along Y-axis), (C) Pressure distribution [Pa], (D) Oxygen distribution

The slice plot presents a series of cross-sectional views that illustrate variations in fluid velocity across the ZX-plane and distributed along the Y-axis. Three equidistant, vertical slices spanning the diameter of the bioreactor were analysed. The results show that fluid velocity peaks near the inlet flow channels and progressively diminishes towards the central region of the bioreactor. This gradient is attributed to the flow dissipative effects as it spreads throughout the volume, ensuring that also those regions distant from the inlets have some level of mixing (Figure 5B).

#### 3.1.2 Pressure Distribution

The pressure distribution analysis highlights the spatial variation of pressure within the bioreactor. The graphical representation, complemented by a color-coded legend (from blue to dark red, as pressure increases), shows that the highest pressure is located at the inlet flow channels. Beyond these regions, the pressure is uniform throughout the vessel, crucial for maintaining stable operating conditions and ensuring the integrity of the biological processes occurring within the bioreactor, as shown in Figure 5(C).

#### 3.1.3 Oxygen Distribution

Oxygen distribution within the BioAxFlow bioreactor was simulated using 3D computational models. The results indicate an average oxygen concentration of 185 uM in the vessel, with a gradient ranging from 220 uM near the air-liquid interface (dark red in Figure 5) to 130 uM near the walls (lighter red in Figure 5D).

### 3.2 Analysis of cell adhesion to bone-mimicking scaffolds using the BioAxFlow cell culture system

This study explored the BioAxFlow (BAF) bioreactor’s potential to enhance cell colonization on biomimetic scaffolds for tissue engineering applications. Specifically, we assessed the adhesion of osteoblast-like SAOS-2 cells to polylactic acid (PLA) bone inorganic matrix-mimicking scaffolds after 24 hours of culture in the BAF bioreactor at two initial cell concentrations: 150,000 cells ml^-1^ (Figure 6B) and 50,000 cells ml^-1^ (Figure 6C). Dynamic cultures were conducted using two BAF bioreactors in two sizes (4 cm and 8 cm in diameter, respectively), while static cultures were performed in standard plastics multi-well plates. Cell adhesion was quantified using the Trypan Blue exclusion assay, which measured the number of nucleated and viable cells after detachment from the scaffolds used. Preliminary experiments were conducted to indirectly demonstrate the BAF bioreactor’s capability to maintain cells in suspension. These experiments evaluated the adhesion of osteoblastic SAOS-2 cells to PLA scaffolds at an initial concentration of 150,000 cells ml^-1^ over different time periods (2h, 4h, 6h, and 24h). The results (Figure 6A) showed that after 2h (blue bar) around 100,000 cells adhered to the scaffold, after 4h (orange bar) cellular adhesion increased to around 150’000 cells, suggesting progressive attachment to the scaffold confirmed at 6h post seeding (grey bar), when a significant increase in cellular adhesion was observed, with over 250,000 cells adhering to the scaffold. Finally at 24 hours (yellow bar), cell adhesion peaked, with approximately 400,000 adherent cells, confirming the time-dependent attachment to the scaffolds as hypothesized. Noticeably, this continuous increase in the number of adherent cells during the first 24 hours of culture, indirectly confirms the ability of the BAF bioreactor to effectively maintain cells in suspension. Indeed, by ensuring that cells remain suspended in the culture medium, the bioreactor facilitates a continuous interaction with the scaffold, thereby enhancing their colonization over time. Following these preliminary findings, we evaluated cell adhesion under dynamic and static conditions (Figure 6B). When an initial concentration of 150,000 cells ml^-1^ was seeded, the dynamic culture (using the 4 cm BAF bioreactor) resulted in approximately 290’000 adherent cells on average, representing a statistically significant increase compared to static culture conditions (where around 101’000 adherent cells were found, on average, per scaffold). This finding indicates a 2.8-fold increase in cellular adhesion onto the PLA-scaffolds when cultured within the BAF bioreactor instead that using static multi-well plates. The larger 8 cm BAF bioreactor yielded even higher cellular adhesion values, with an average of 380,000 cells per scaffold—a 3.8-fold statistically significant increase compared to static conditions. These results indicate that the continuous flow environment in the BAF bioreactor enhances cellular colonization by improving cell suspension and exposure to nutrients and growth factors. To further analyse the impact of initial cellular concentration on scaffolds’ colonization, SAOS-2 cells were also seeded at a lower concentration of 50,000 cells ml^-1^ and cultured in the 8 cm BAF bioreactor (Figure 6C). Under these conditions, dynamic culture resulted in approximately 176,000 cells adhered per scaffold, a statistically significant improvement compared to static culture, which yielded only 33,000 adhered cells per scaffold. Despite the lower initial seeding concentration, the fold increase in cellular adhesion under dynamic conditions within BAF (5.4-fold) was significantly higher than that observed when using the 150,000 cells ml^-1^ cellular concentration for seeding (3.8-fold). These findings confirm that the dynamic environment provided by the BAF bioreactor significantly improves cellular colonization onto biomimetic scaffolds, making it a promising tool for tissue engineering applications.

**Figure 6.**
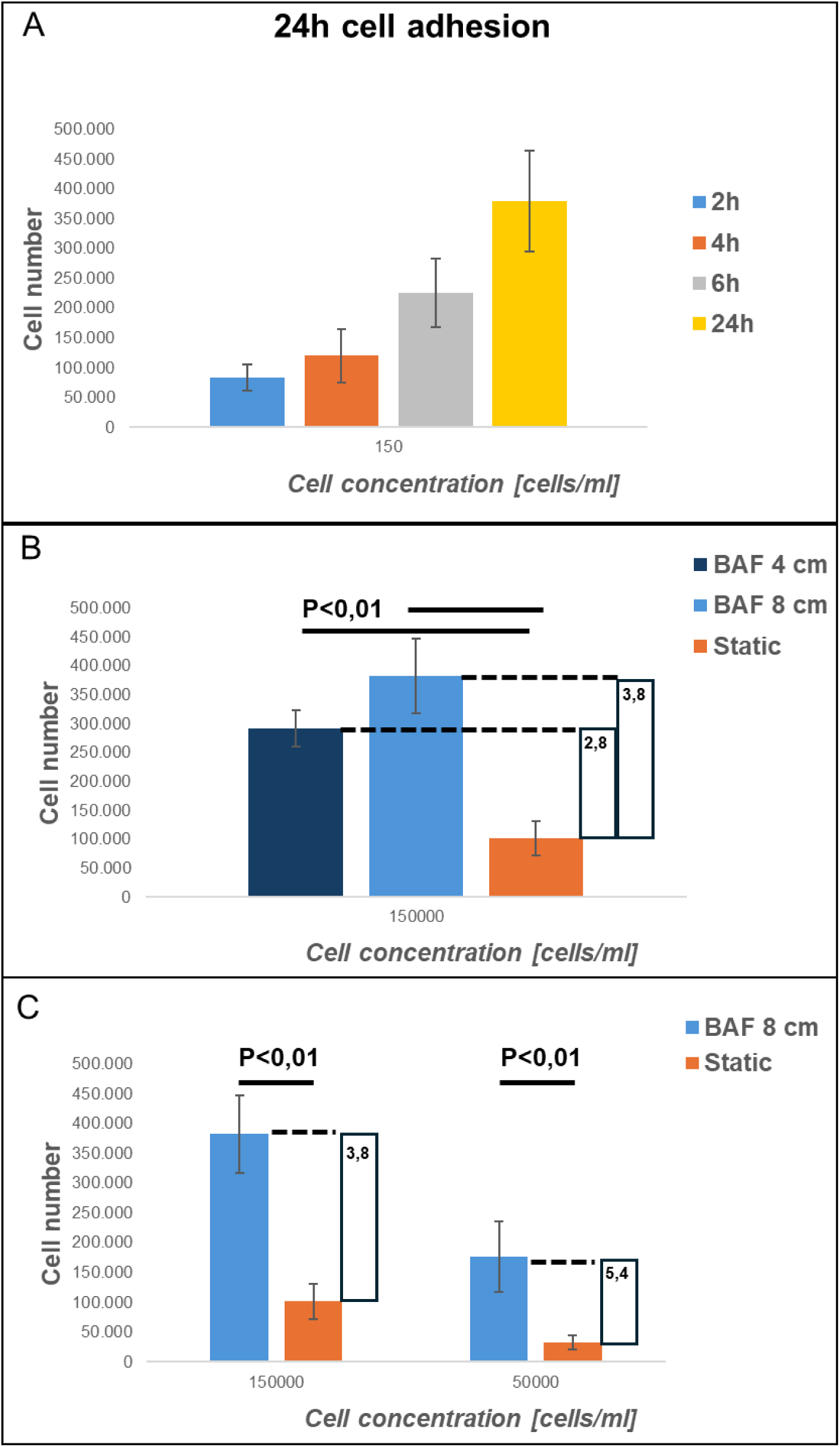
Analysis of cell adhesion at 24h of cells cultured on PLA scaffolds under different experimental conditions: using BioAxFlow (BAF, in light blue) versus multi-well plates (static conditions, in orange). Data are shown as mean SD. n=6, for each condition (BAF 4 cm, BAF 8 cm, static conditions).

### 3.3 Analysis of cell growth on bone-mimicking scaffolds within the BioAxFlow cell culture system

The results of the cell adhesion analysis demonstrate that the BAF bioreactor significantly enhances cellular adhesion, achieving approximately a five-fold increase compared to static conditions, depending on the bioreactor’s size. To further evaluate the impact of the usage of BAF on cell growth, SAOS-2 cells were grown on bone-like scaffolds, allowing their adhesion for 24 hours inside the BAF bioreactor and subsequently continued to be cultured in the same bioreactor for up to 10 days. Cell growth was quantified at days 1, 4, 7, and 10 by counting the number of viable cells on the scaffolds using the Trypan Blue exclusion assay. The growth curves were obtained using two initial SAOS-2 cellular concentrations (150,000 cells ml^-1^ and 50,000 cells ml^-1^), and the results of the experiments conducted under dynamic (BAF bioreactor) or static conditions are shown in Figure 7. When an initial concentration of 150,000 cells ml^-1^ was used (Figure 7A), under dynamic conditions in the BAF bioreactor, the number of counted cells increased steadily from day 1, reaching approximately 4.0 million cells by day 7. By contrast, in static conditions, cellular proliferation was much slower, reaching only about 1.3 million cells by day 7. By the end of the experiment, cellular proliferation, when using BAF for culture, was approximately three times more than the one when using multi-well plates. This demonstrates the significant advantage derived from the usage of dynamic culture conditions in supporting rapid and efficient cell growth over a shorter timeframe. However, at the higher seeding cellular concentration of 150000 cells ml^-1^, some experiments revealed the presence of non-viable cells at day 7, particularly in static conditions. This was likely due to nutrient depletion, waste accumulation, and reduced oxygen availability in the scaffolds’ microenvironment, impairing cell viability. To avoid confounding effects associated with compromised cellular vitality, the experiment at this concentration was terminated on Day 7. To investigate cell growth over an extended period, cultures were set up with a lower seeding concentration of 50,000 cells ml^-1^ (Figure 7B). Under dynamic conditions, the number of counted cells increased significantly, reaching approximately 4.2 million cells by day 10. This consistent and robust proliferation demonstrates the ability of the BAF bioreactor to sustain cellular growth over a longer timeframe. In contrast, static culture conditions supported much slower growth, with the cell number reaching a maximal value of approximately 1.1 million by day 10. Thus, the BAF bioreactor supported nearly four times greater number of cells compared to static conditions, confirming its ability to sustain high proliferation rates even with lower initial cell seeding. Statistical analysis via two-way ANOVA confirmed that, for both seeding concentrations, cell growth under dynamic conditions was significantly higher than under static conditions (p < 0.01)). The superior performance of the BAF bioreactor is attributed to its continuous flow system, which ensures optimal nutrient distribution, oxygen supply, and waste removal, hence preventing to reach the plateau and decline phases of the cellular growth curve observed in static culture conditions, where limited diffusion likely restricts cell viability and proliferation. These results highlight the efficacy of the BAF bioreactor in supporting enhanced cell growth and its potential as a valuable tool for tissue engineering applications.

**Fig. 7.**
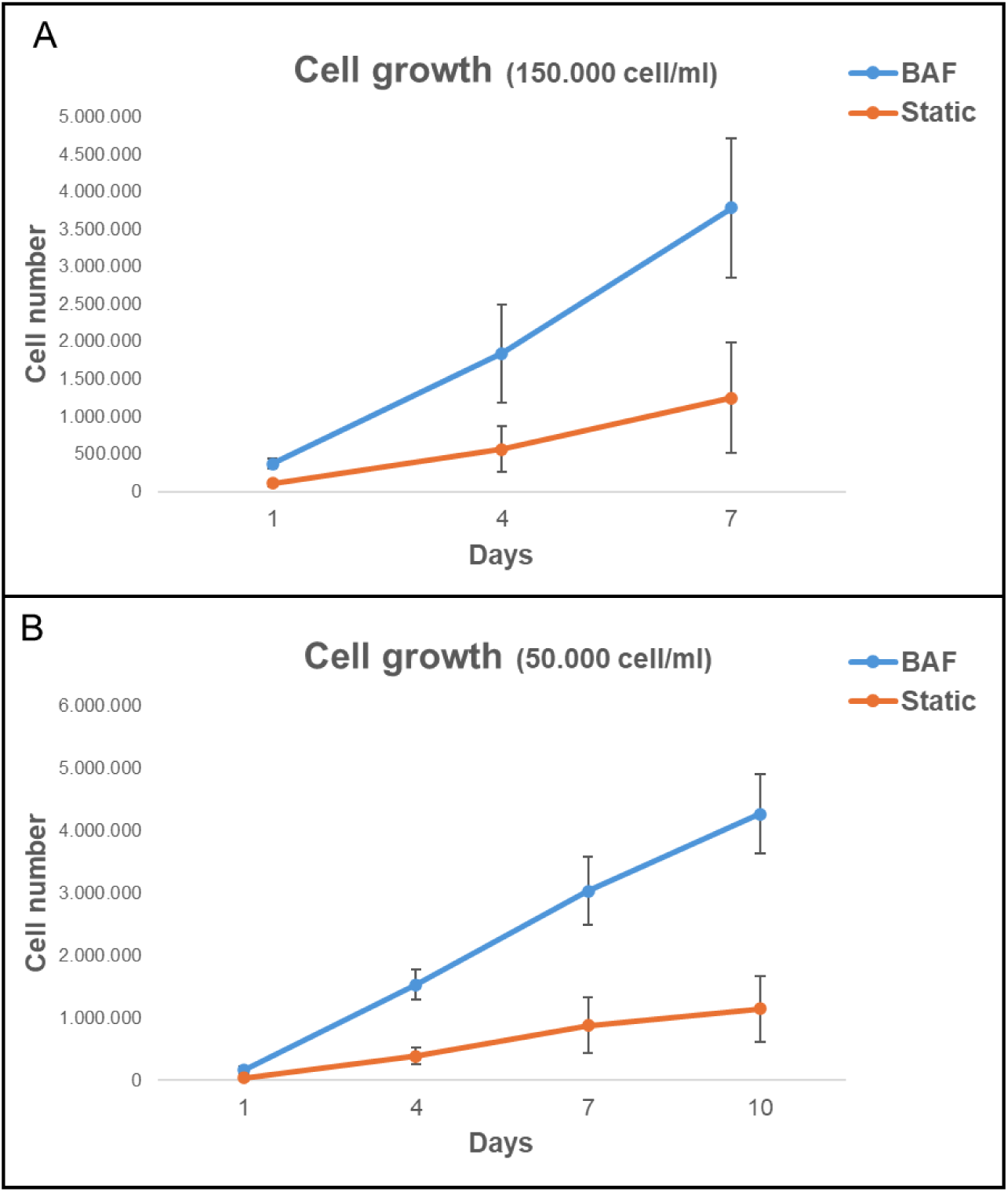
Analysis of cell growth on PLA scaffolds under different experimental conditions in the bioreactor BAF versus static conditions. The growth curves were obtained using two initial SAOS-2 cell concentrations: (A) 150,000 cells ml^-1^ and (b) 50,000 cells ml^-1^. Data are shown as mean SD. n=6 for each experimental condition.

### 3.4 Effect of BioAxFlow on the transitions from Dynamic to Static Culture on Cell Proliferation

The experiments described above suggest that using the BAF bioreactor consistently supports a higher cell proliferation rate compared to cultivating in static conditions. To determine whether this effect was due only to differences in initial cell adhesion or to better culture conditions within the bioreactor, scaffolds were subjected to two experimental setups: i) they were cultured in static conditions for 24 hours (for cell seeding) and then transferred to the bioreactor for dynamic culture up to 10 days (Static to BAF); ii) Scaffolds were cultured in the bioreactor for 24 hours (for cell seeding) before being transferred to static culture conditions for up to 10 days (BAF to Static). The results, shown in Figure 8, demonstrate that scaffolds shifted from static to dynamic conditions (Static to BAF) exhibited a statistically significant increase in cell proliferation rates compared to purely static culture. By day 10, cell counts reached approximately 2.8 million, confirming the bioreactor’s positive impact on both cell adhesion and proliferation. In contrast, scaffolds transferred from dynamic to static conditions after 24 hours of seeding (BAF to Static) showed a consistent slowdown in cell proliferation rates, as expected. Before being moved to static conditions, the cell adhesion achieved with the BAF bioreactor was higher than that achieved with static culture systems. However, the growth rate slowed significantly once the scaffolds were removed from the dynamic environment and by day 10, cell counts reached approximately 1.7 million, indicating the reduced benefits of the bioreactor’s culture conditions once removed.

**Figure 8.**
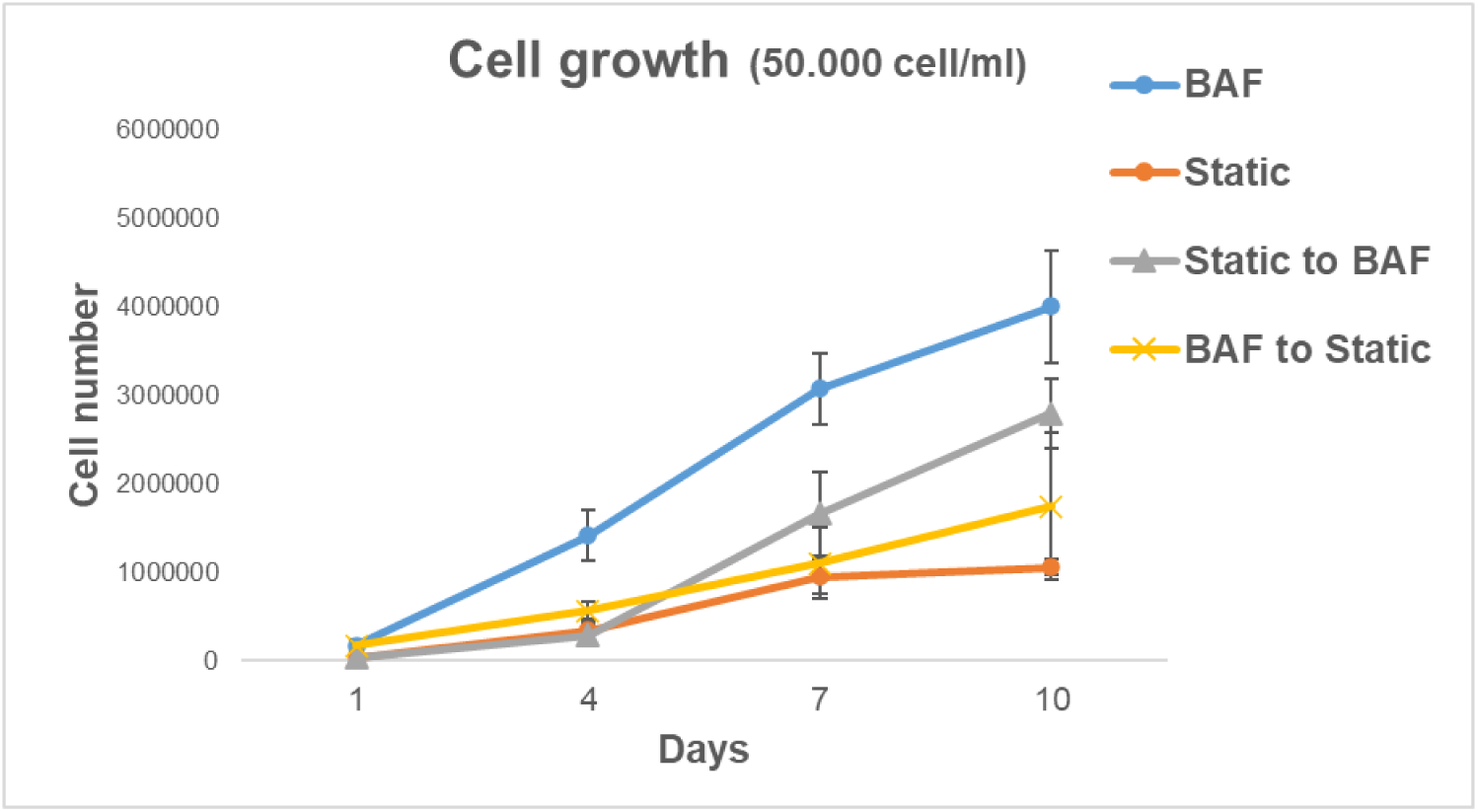
Analysis of cell growth on scaffold after the shift from static to dynamic conditions (Static to BAF) and from dynamic to static conditions (BAF to Static). The growth curves were obtained using the same SAOS-2 cellular concentration of 50,000 cells ml^-1^ at seeding. Data are shown as mean SD. n=6 for each experimental condition.

### 3.5 Analysis of cell distribution across the different surfaces of the scaffolds

The cellular colonization was further studied in terms of cellular distribution across the different surfaces of the scaffolds, comparing the dynamic 3D culture in the bioreactor to traditional static culture methods. This was achieved through fluorescent dye staining (propidium iodide) of the cell nuclei and subsequent visualization under confocal microscopy. Figure 9 reports the images of the entire surface of scaffolds, acquired at 4x magnification. The static culture conditions do not allow for homogeneous colonization of the scaffolds, in contrast to what is observed in the dynamic culture conditions achieved using BAF. Under static culture conditions, cells primarily adhere to surface A, which is the upper surface where the cell suspension is directly deposited during seeding. In contrast, surface B, oriented towards the bottom of the plate, shows limited cell colonization. Conversely, the dynamic 3D cell culture conditions, achieved using BAF, led to the colonization of both surfaces of each scaffold. This is attributed to the fluid dynamics generated within the bioreactor, which keeps the cells in suspension, gently mixes them and permits a homogeneous mass transfer.

**Figure 9.**
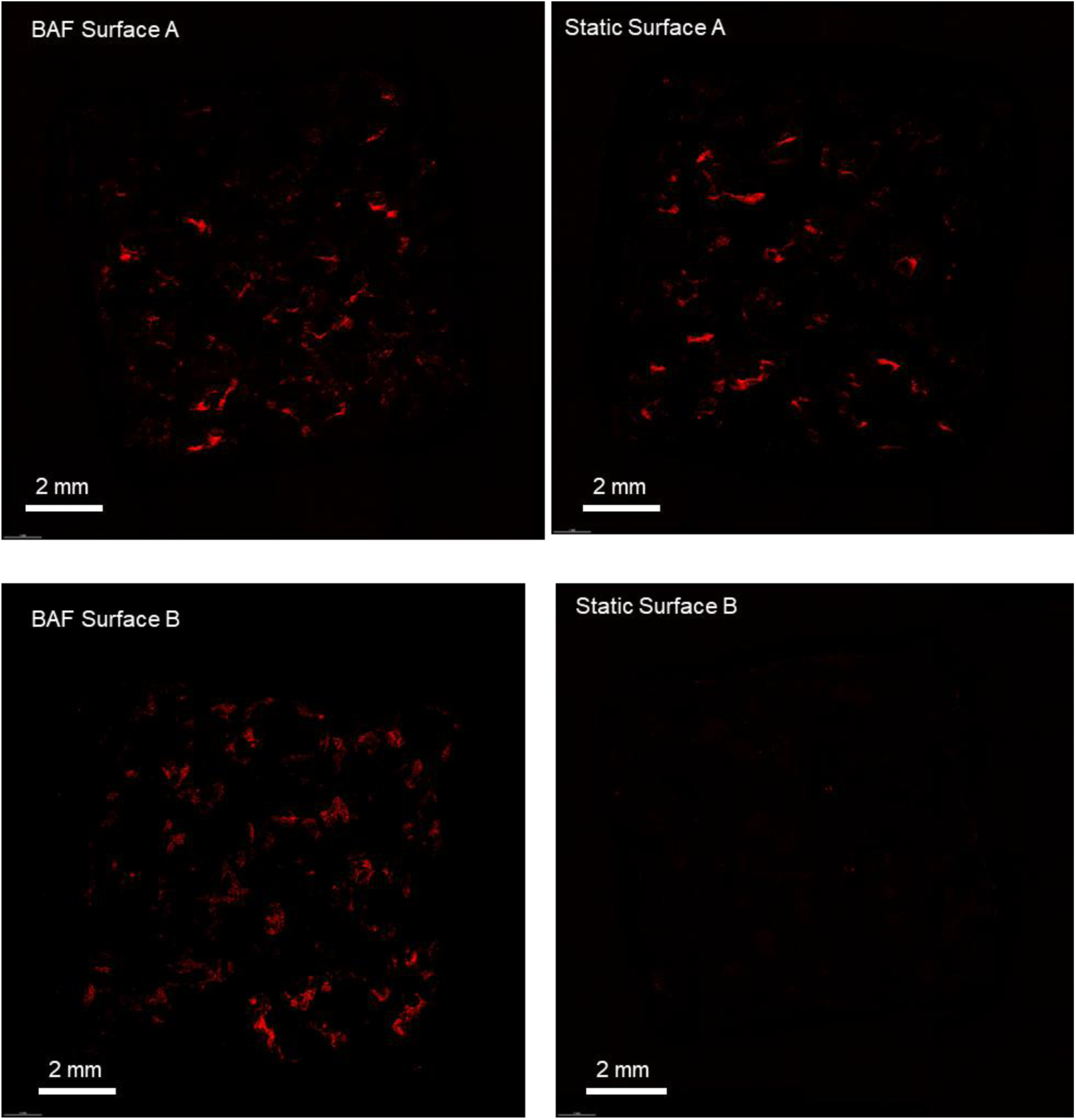
Confocal microscopy analysis of SAOS-2 cells cultured on scaffolds, under dynamic and static conditions, revealed by fluorescent dye staining (propidium iodide) of the cell nuclei. Photographs were taken at a magnification of 4X.

### 3.6 Analysis of mRNA expression in SAOS-2 cells grown within the BioAxFlow cell culture system

#### 3.6.1 Expression of constitutive and osteogenic mRNA markers

The above-described results demonstrate that the BAF bioreactor significantly enhances biomaterial-made scaffolds colonization, promoting a uniform cell distribution across all surfaces. Additionally, the fluid dynamics generated within the bioreactor improves cell growth compared to static conditions. To further confirm the bioreactor’s biocompatibility and evaluate its osteoconductive potential, the expression of mRNAs associated with osteogenic differentiation and essential for biosynthetic activity, was examined. Specifically, the expression levels of key constitutive genes—Ki67, and RPL34—as well as osteogenic markers such as osteopontin (OPN), alkaline phosphatase (ALP), RUNX2, and osteocalcin (OCL) were studied by qRT-PCR assay. Ki67 is a proliferation marker, involved in cell cycle; and RPL34, a ribosomal protein, is essential during the translation process. These transcripts, highly expressed in SAOS-2 cells, are recognized as reliable indicators of correct cell functions. Their expression levels were assessed at day 1, 4 and 7 post seeding within the bioreactor and found to be comparable to those under static conditions (Figure 10A). Noticeably, the bioreactor’s ability to maintain the osteogenic commitment of SAOS-2 cells was also validated. This commitment is a key indicator of the cells’ capacity to preserve their osteogenic potential and osteoconductive properties. Osteogenesis involves a sequence of events across three stages: (1) proliferation, (2) extracellular matrix (ECM) deposition and maturation, and (3) mineralization of the bone ECM (30). These phases are driven by transcription factors like RUNX2 and the activation of osteoblast-specific genes, including ALP, OCL, and OPN. ALP, an early differentiation marker, is expressed towards the end of the proliferative phase and during ECM maturation, while OPN and OCL— specific markers of the mineralization phase—are associated with advanced osteogenesis (31). The qRT-PCR analysis of osteogenic markers, conducted at day 1, 4 and 7 post cellular seeding on PLA scaffolds within the bioreactor, showed no statistically significant differences in the expression of these markers compared to the static control (Figure 10B). These findings confirm that the dynamic environment provided by the BAF bioreactor preserves the osteogenic phenotype of SAOS-2 cells, ensuring their ability to sustain differentiation under dynamic culture conditions.

**Figure 10.**
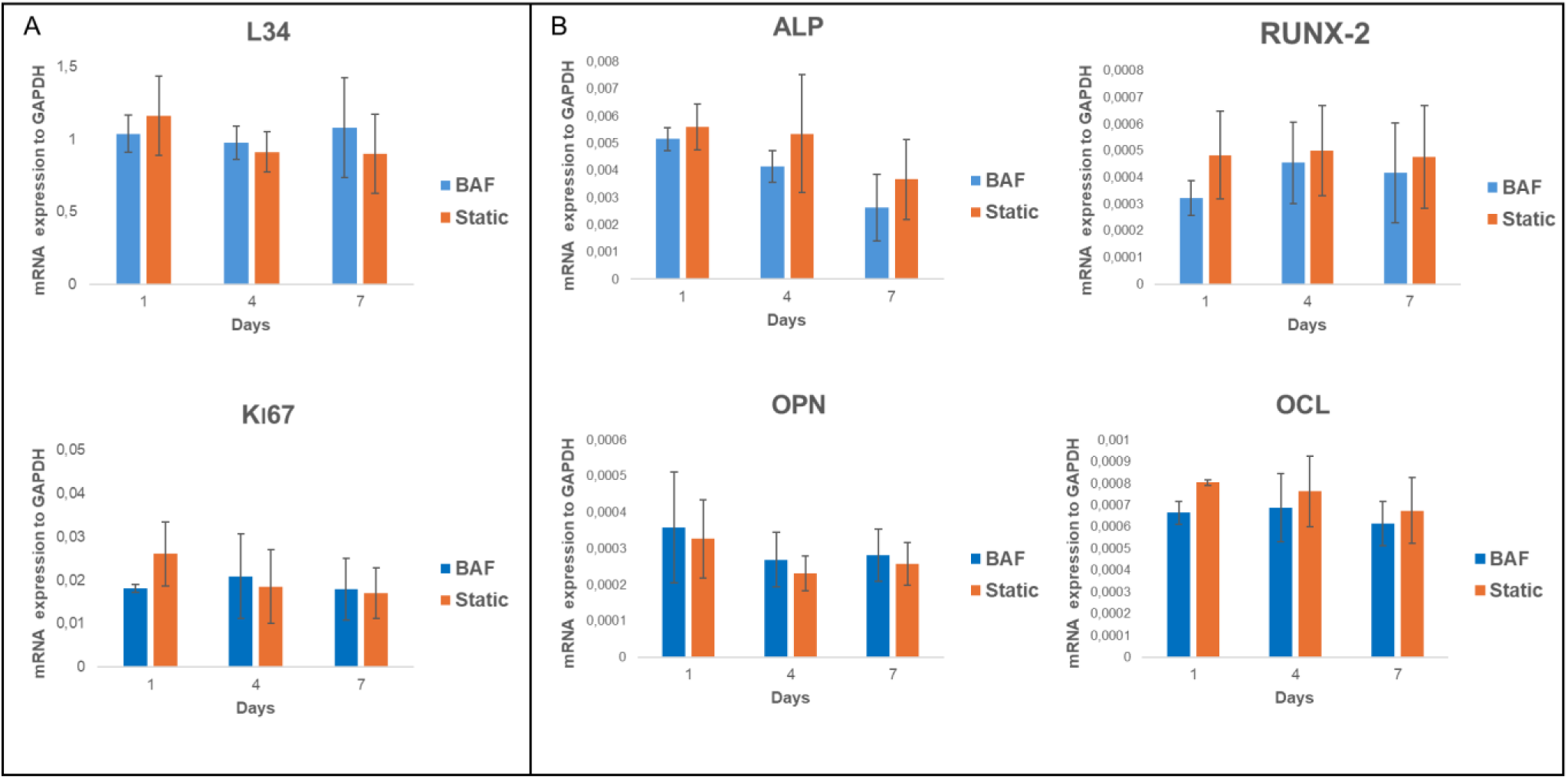
qRT-PCR analysis of the expression of key constitutive (L34, Ki67) and differentiation markers (RUNX2, AP, OPN, and OCL) in SAOS-2 cells cultured for up to 7 days on PLA scaffolds under dynamic BAF bioreactor conditions (BAF) versus static conditions (Static), normalized to the expression of the constitutive gene GAPDH. Data are shown as mean SD. n=3 for each gene analysed, in both experimental conditions (BAF VS Static conditions)

#### 3.6.2 Expression of tumour-related mRNA markers

The SAOS-2 cell line is a widely used model in osteosarcoma research due to its osteoblastic characteristics and ability to mimic bone biology and cancer pathology. These cells exhibit rapid proliferation, resistance to apoptosis, express bone-specific markers and secrete VEGF, which promotes the formation of new blood vessels, a key step in tumour growth and metastasis. To investigate the impact of dynamic culture conditions on the tumorigenic properties of SAOS-2 cells, we analysed the mRNA expression of VEGF, BAX, and BCL2 genes. These genes expression was assessed by qPCR in cells cultured on PLA scaffolds under dynamic conditions (using BAF bioreactor) and static conditions over 1, 4, and 7 days. The results, reported in figure 11, showed no significant changes in VEGF, BAX, or BCL2 expression between dynamic and static conditions, indicating that dynamic culture does not alter key properties related to cancer progression such as angiogenesis or apoptosis regulation. These findings suggest that the BAF bioreactor provides a stable, biocompatible environment that maintains the cancer-like characteristics of SAOS-2 cells, making it a useful platform for in vitro studies on osteosarcoma therapies, targeting angiogenic and apoptotic pathways.

**Figure 11.**
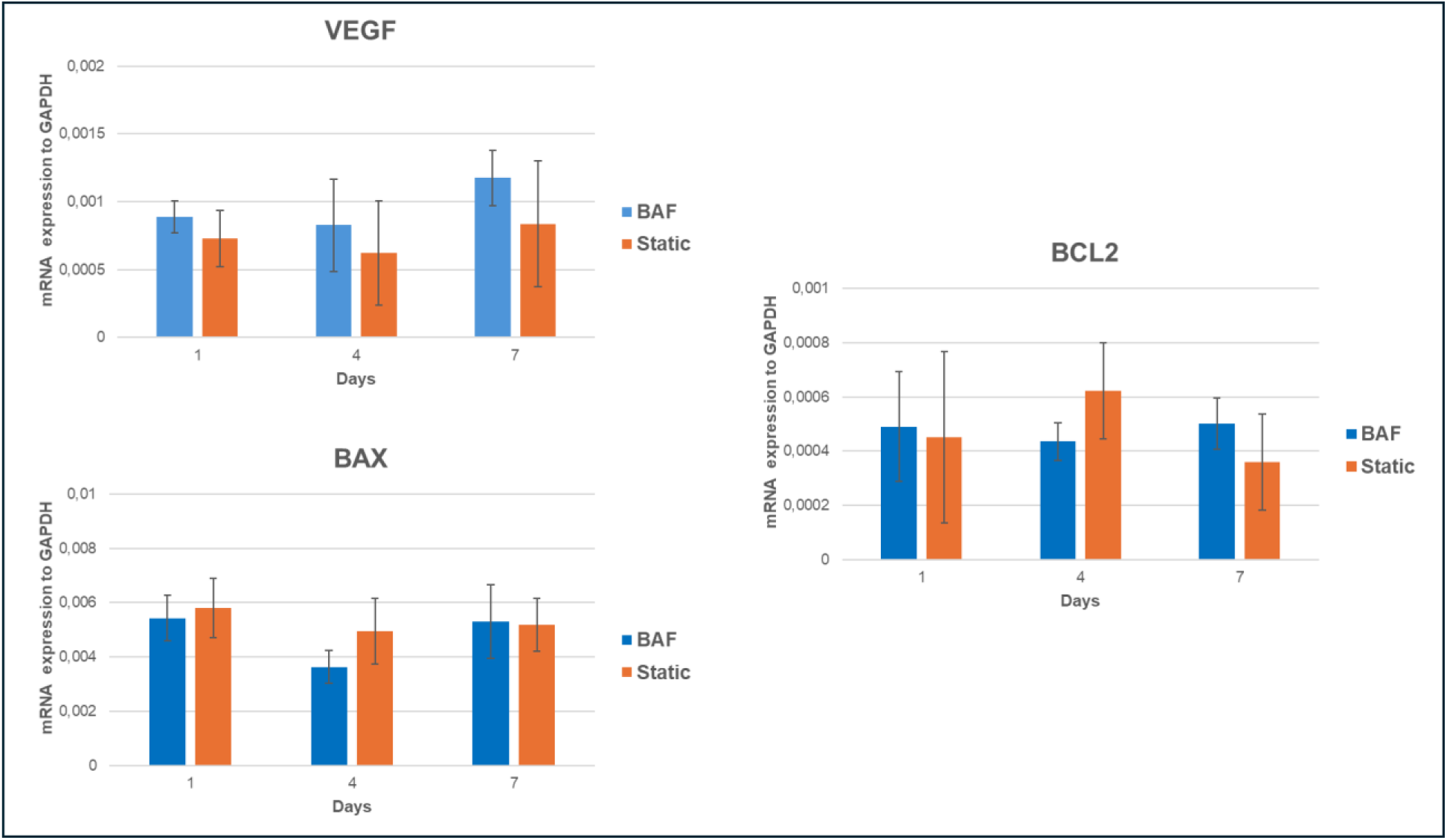
qRT-PCR analysis of VEGF, BAX, and BCL2 mRNA expression in SAOS-2 cells cultured onto PLA scaffolds under dynamic conditions compared to static ones, normalized to the expression of the constitutive gene GAPDH. Data are shown as mean SD. n=3 per experimental condition, at each assessment timing (1, 4 and 7 days)

## 4 Discussion

In this study, we assessed the potential of the BioAxFlow (BAF) bioreactor to enhance the colonization, growth, and to maintain the osteogenic potential of SAOS-2 osteosarcoma cells cultured onto bone-mimicking scaffolds made of polylactic acid (PLA). The results demonstrated that dynamic culture conditions within the BAF bioreactor significantly improved cell adhesion and growth compared to traditional static conditions, highlighting the bioreactor’s potential to be used as a robust platform for tissue engineering applications, particularly in bone regeneration and cancer research.

Our results revealed that dynamic culture in the BAF bioreactor led to a substantial increase in cell adhesion compared to static conditions. At a seeding cellular concentration of 150,000 cells ml^-1^, the 4 cm BAF bioreactor promoted a significant increase in adherent cells, that was enhanced when the 8 cm bioreactor was used. Notably, even at a lower seeding density of 50,000 cells ml^-1^, dynamic conditions in the larger bioreactor (8 cm), resulted in a further improvement in the average cellular adhesion. This underscores the BAF system’s ability to maintain effective cell-scaffold interactions despite reduced cell availability, consistent with previous studies that have demonstrated the benefits of dynamic fluid-based culture systems for promoting efficient cell seeding and biomaterial colonization (32).The fluid dynamics within the bioreactor, as highlighted by the COMSOL simulations, ensures uniformity in key parameters such as velocity, pressure, and gas distributions. Specifically, the simulations demonstrate the capability to achieve a homogeneous velocity and oxygen distribution throughout the system, which is critical for a homogeneous distribution of cells and nutrients throughout the entire volume of the scaffolds. The enhanced adhesion observed is likely due to the continuous fluid flow within the bioreactor, which improves nutrient and growth factor exchange while mimicking physiological environments more closely than static culture. This promotes greater cellular interaction with biomaterial surfaces, making the BAF bioreactor able to support an efficient scaffold seeding and better nutrient exposure. Furthermore, the improved adhesion at lower cellular concentrations at seeding (50,000 cells ml^-1^) demonstrates the bioreactor’s efficiency and scalability. The bioreactor’s advantage extended beyond initial cell adhesion, demonstrating superior performance in supporting sustained cell proliferation over time. At an initial seeding cellular concentration of 150,000 cells ml^-1^, dynamic culture resulted in a 3-fold increase in number of cells by day 7 compared to static culture. Importantly, the experiment revealed reduced viability in static conditions after day 7, likely due to nutrient and oxygen depletion, alongside waste accumulation, within the scaffolds’ microenvironment. These findings align with previous reports emphasizing the challenges of maintaining cellular vitality in static cultures, particularly at higher seeding cellular densities (33, 34). At a lower seeding cellular concentration of 50,000 cells ml^-1^, the BAF bioreactor enabled consistent cell growth, achieving a nearly 4-fold higher number of cells compared to static culture by day 10. This sustained proliferation highlights the capacity of the bioreactor to maintain optimal growth conditions over extended periods. By ensuring continuous nutrient delivery and waste removal, the BAF bioreactor mitigates the diffusion limitations inherent to static cultures, supporting long-term viability and robust cellular expansion. This is consistent with findings from similar studies that show enhanced cell proliferation in bioreactors due to continuous nutrient supply, prevention of nutrient depletion, and waste accumulation that typically limit cell growth in static systems (35). To assess whether the beneficial effects on cultured cells are merely linked to higher initial cellular adhesion to the scaffolds or to better culture conditions achieved within the bioreactor, we used an approach consisting in culturing the scaffolds in static conditions for 24 hours and then transferring them to the bioreactor, while other scaffolds, originally cultured in the bioreactor for 24 hours, were shifted to static culture conditions. Scaffolds shifted from static to dynamic conditions (Static to BAF, in Figure 8) exhibited a significant increase in cellular proliferation comparted to static culture. Conversely, scaffolds moved from dynamic to static conditions (BAF to Static, in Figure 8) displayed slower growth, indicating reduced proliferation after removal from the bioreactor’s dynamic environment. These results highlight the critical role of the bioreactor’s dynamic culture environment in fostering robust and sustained cell proliferation. The continuous flow of culture medium achieved in the bioreactor’s culture chamber, being steadily gentle thanks to the pairing to a peristaltic pump, ensures an optimal balance between nutrient supply, oxygenation, and waste removal, preventing the plateau in the growth curve observed in static cultures to occur (see Figure 8). The recovery of cellular growth rate observed when scaffolds were shifted from static to dynamic conditions underscores the bioreactor’s effectiveness in enhancing proliferation even after initial suboptimal conditions. Conversely, the reduction in the growth rate observed when scaffolds were transitioned from dynamic to static conditions highlights the importance of sustained dynamic culture for maintaining cell proliferation and viability over time at optimal rates. By offering efficient scaffold seeding, robust cell expansion, and long-term viability, the bioreactor addresses critical challenges in scalable tissue engineering strategies. The BAF bioreactor not only supports rapid and efficient cell adhesion to the bone-mimicking scaffolds specifically designed for this study but also provides optimal conditions for sustained cell growth.

A crucial aspect in tissue engineering is the ability to achieve homogeneous cellular distribution across the entire scaffold surface. When cultured using traditional static conditions, SAOS-2 cells’ distribution was limited to the surface directly exposed to the cell suspension, with minimal colonization observed on the opposite surface. In contrast, dynamic conditions facilitated uniform cell distribution across both scaffold surfaces, as demonstrated by fluorescent staining by confocal microscopy that can be attributed to the fluid dynamics within the BAF bioreactor, which helps in maintaining cells in suspension, thereby promoting even seeding and preventing cells from settling unevenly. This finding underscores the importance of dynamic culture systems for achieving effective cellular colonization in three-dimensional scaffolds, which is essential for developing functional tissues for regenerative medicine.

In addition to evaluating the bioreactor’s impact on cellular adhesion and growth, we also investigated the effect of dynamic culture on the osteogenic potential of SAOS-2 cells. The expression of key osteogenic markers, such as RUNX2, ALP, OPN, and OCL, was assessed over time under both dynamic and static culture conditions. Our data revealed that the expression of these osteogenic markers remained comparable between dynamic and static cultures, indicating that the dynamic environment provided by the BAF bioreactor did not compromise the osteogenic commitment of SAOS-2 cells. This result is consistent with previous studies, which suggest that bioreactors can support the osteogenic differentiation of osteoblast-like cells by maintaining physiological conditions (36). Therefore, the BAF bioreactor offers a biocompatible environment for long-term culture of SAOS-2 cells while preserving their osteogenic potential.

A key aspect of our study was evaluating the impact of dynamic culture conditions on the tumour phenotype of SAOS-2 cells, which are commonly used as a model for osteosarcoma. We analysed the expression of tumour-related markers, including VEGF (a key pro-angiogenic factor), and apoptotic genes, BAX (pro-apoptotic) and BCL2 (anti-apoptotic), to assess whether dynamic culture within the BAF bioreactor would alter the tumorigenic potential of SAOS-2 cells. Our results showed that there were no significant changes in the expression of these markers under dynamic conditions, indicating that the bioreactor did not alter the tumorigenic potential of SAOS-2 cell, being ineffective in altering their angiogenic or apoptotic regulation. Specifically, VEGF expression remained stable, suggesting that dynamic conditions did not exacerbate the pro-angiogenic phenotype of the cells. Similarly, the balance between BAX and BCL2 was maintained, supporting the conclusion that the BAF bioreactor does not induce significant apoptotic stress. These findings suggest that the BAF bioreactor can be used as an effective platform for studying the tumorigenic characteristics of osteosarcoma cells in vitro without altering their tumorigenic potential. The ability to culture SAOS-2 cells under dynamic conditions while preserving their tumour phenotype is particularly valuable for both drug screening and testing of therapies targeting angiogenesis and apoptosis.

Bone tissue engineering requires effective colonization of biomimetic scaffolds to establish functional constructs that integrate seamlessly with host tissue. The BAF bioreactor’s controlled dynamic environment promotes rapid cell colonization, uniform distribution across bone-mimicking scaffolds, and high cellular viability. Improvements at lower seeding concentrations highlight its potential for cost-effective and efficient cell culture strategies, minimizing resource demands while maintaining high-quality outcomes. This advantage is particularly beneficial for large-scale applications and the development of complex, multilayered constructs. Future studies should explore varying flow rates and scaffold geometries to optimize performance and scalability for clinical applications. Additionally, incorporating co-culture systems and diverse cell types would provide valuable insights into the bioreactor’s versatility and broader applicability.

## 5 Conclusion

In conclusion, the BioAxFlow (BAF) bioreactor represents a powerful tool for advancing tissue engineering and cancer research. This study highlights its ability to significantly enhance cellular adhesion, proliferation, and uniform scaffold colonization compared to traditional static culture methods. The flow generated within the culture chamber ensures continuous movement of the medium, promoting consistent cellular exposure to nutrients and growth factors, which facilitates a more efficient nutrient exchange and cellular interaction with the scaffolds, closely mimicking the conditions observed in vivo. By providing a dynamic culture environment that mimics physiological conditions, the BAF bioreactor overcomes the limitations of diffusion-based transport, such as nutrient depletion and waste accumulation, enabling rapid and sustained cell growth while maintaining cell viability and cell commitment. Notably, the BAF bioreactor offers a biocompatible and stable platform suitable for studying osteosarcoma models, as it preserves the tumorigenic properties of SAOS-2 cells while still enabling researchers to design specific in vitro experiments. This makes the BAF bioreactor particularly valuable for exploring cancer therapies targeting angiogenesis and apoptosis, e.g. in a patient-specific manner. Furthermore, the system’s versatility extends beyond bone tissue engineering and osteosarcoma research, with potential applications in studying a wide range of tumour types and conducting cellular differentiation studies. Its adaptability underscores its capacity to support innovative approaches in both regenerative medicine and oncological studies. These findings pave the way for future advancements in scaffold-based tissue engineering and in vitro cancer research facilitating the development of novel therapeutic strategies and demonstrating the potential of bioreactor technologies to revolutionize how current pre-clinical studies are conducted.

## Acknowledgements

This work has been funded by the Following project:

-European Commission project OSTEONET - HORIZON-MSCA-2021-SE-01 GA n. 101086329.

-Lazio Region project 3DCREATOR Programma FESR Lazio 2021-2027 “Riposizionamento Competitivo” CUP “B53D23034400001.

- MUR DSB.AD006.371.010 / Invecchiamento attivo e in salute (FOE 2022) IFT

## Author Contributions

M.L., A.L. and G.F.D.L contributed equally to this work. G.G and F.L. did the experimental validation and writing, LF contributed to writing and reviewing, M.M, M.S. M.P and F.L. developed the technology of BioAxFlow based on GFDL owned IPRs.

## References

1. Bolamperti S, Villa I, Rubinacci A. Bone remodeling: an operational process ensuring survival and bone mechanical competence. Bone Res. 2022;10(1):48.

2. Lopez-Otin C, Blasco MA, Partridge L, Serrano M, Kroemer G. The hallmarks of aging. Cell. 2013;153(6):1194–217.

3. Ensrud KE, Crandall CJ. Osteoporosis. Ann Intern Med. 2017;167(3):ITC17–ITC32.

4. Keaveny TM, Hayes WC. A 20-year perspective on the mechanical properties of trabecular bone. J Biomech Eng. 1993;115(4B):534–42.

5. Zebaze RM, Ghasem-Zadeh A, Bohte A, Iuliano-Burns S, Mirams M, Price RI, et al. Intracortical remodelling and porosity in the distal radius and post-mortem femurs of women: a cross-sectional study. Lancet. 2010;375(9727):1729–36.

6. McBroom RJ, Hayes WC, Edwards WT, Goldberg RP, White AA, 3rd. Prediction of vertebral body compressive fracture using quantitative computed tomography. J Bone Joint Surg Am. 1985;67(8):1206–14.

7. Mosekilde L, Mosekilde L. Normal vertebral body size and compressive strength: relations to age and to vertebral and iliac trabecular bone compressive strength. Bone. 1986;7(3):207–12.

8. Liu XS, Sajda P, Saha PK, Wehrli FW, Guo XE. Quantification of the roles of trabecular microarchitecture and trabecular type in determining the elastic modulus of human trabecular bone. J Bone Miner Res. 2006;21(10):1608–17.

9. Pearce AI, Richards RG, Milz S, Schneider E, Pearce SG. Animal models for implant biomaterial research in bone: a review. Eur Cell Mater. 2007;13:1–10.

10. Mhanna R, Hasan A. Introduction to Tissue Engineering. Tissue Engineering for Artificial Organs2017. p. 1–34.

11. Bouet G, Marchat D, Cruel M, Malaval L, Vico L. In vitro three-dimensional bone tissue models: from cells to controlled and dynamic environment. Tissue Eng Part B Rev. 2015;21(1):133–56.

12. Kumari S, Katiyar S, Anand A, Singh D, Singh BN, Mallick SP, et al. Design strategies for composite matrix and multifunctional polymeric scaffolds with enhanced bioactivity for bone tissue engineering. Front Chem. 2022;10.

13. Lawrence LM, Salary RR, Miller V, Valluri A, Denning KL, Case-Perry S, et al. Osteoregenerative Potential of 3D-Printed Poly epsilon-Caprolactone Tissue Scaffolds In Vitro Using Minimally Manipulative Expansion of Primary Human Bone Marrow Stem Cells. Int J Mol Sci. 2023;24(5).

14. Ledda M, Merco M, Sciortino A, Scatena E, Convertino A, Lisi A, et al. Biological Response to Bioinspired Microporous 3D-Printed Scaffolds for Bone Tissue Engineering. Int J Mol Sci. 2022;23(10).

15. Saldana L, Bensiamar F, Bore A, Vilaboa N. In search of representative models of human bone-forming cells for cytocompatibility studies. Acta Biomater. 2011;7(12):4210–21.

16. Steinerova M, Matejka R, Stepanovska J, Filova E, Stankova L, Rysova M, et al. Human osteoblast-like SAOS-2 cells on submicron-scale fibers coated with nanocrystalline diamond films. Mater Sci Eng C Mater Biol Appl. 2021;121:111792.

17. Fernandes RJ, Harkey MA, Weis M, Askew JW, Eyre DR. The post-translational phenotype of collagen synthesized by SAOS-2 osteosarcoma cells. Bone. 2007;40(5):1343–51.

18. Assi T, Watson S, Samra B, Rassy E, Le Cesne A, Italiano A, et al. Targeting the VEGF Pathway in Osteosarcoma. Cells. 2021;10(5).

19. Pautke C, Schieker M, Tischer T, Kolk A, Neth P, Mutschler W, et al. Characterization of osteosarcoma cell lines MG-63, Saos-2 and U-2 OS in comparison to human osteoblasts. Anticancer Res. 2004;24(6):3743–8.

20. Cimmino A, Fasciglione GF, Gioia M, Marini S, Ciaccio C. Multi-Anticancer Activities of Phytoestrogens in Human Osteosarcoma. Int J Mol Sci. 2023;24(17).

21. Lv T, Jian Z, Li D, Ao R, Zhang X, Yu B. Oxyresveratrol induces apoptosis and inhibits cell viability via inhibition of the STAT3 signaling pathway in Saos-2 cells. Mol Med Rep. 2020;22(6):5191–8.

22. Bušs A, Veģere K, Suleiko A, Vanags J, editors. CFD Analysis of a Stirred Vessel Bioreactor with Double Pitch Blade and Rushton Type Impellers2017.

23. Logan DL. A First Course in the Finite Element Method: Thomson; 2007.

24. Mazzei D, Guzzardi MA, Giusti S, Ahluwalia A. A low shear stress modular bioreactor for connected cell culture under high flow rates. Biotechnol Bioeng. 2010;106(1):127–37.

25. Shapovalov Y, Hoffman D, Zuch D, de Mesy Bentley KL, Eliseev RA. Mitochondrial dysfunction in cancer cells due to aberrant mitochondrial replication. J Biol Chem. 2011;286(25):22331–8.

26. Foy BD, Rotem A, Toner M, Tompkins RG, Yarmush ML. A device to measure the oxygen uptake rate of attached cells: importance in bioartificial organ design. Cell Transplant. 1994;3(6):515–27.

27. Buchwald P. A local glucose-and oxygen concentration-based insulin secretion model for pancreatic islets. Theor Biol Med Model. 2011;8:20.

28. Zenobi E, Merco M, Mochi F, Ruspi J, Pecci R, Marchese R, et al. Tailoring the Microarchitectures of 3D Printed Bone-like Scaffolds for Tissue Engineering Applications. Bioengineering (Basel). 2023;10(5).

29. Livak KJ, Schmittgen TD. Analysis of relative gene expression data using real-time quantitative PCR and the 2(-Delta Delta C(T)) Method. Methods. 2001;25(4):402–8.

30. Maccioni RB, Cambiazo V. Role of microtubule-associated proteins in the control of microtubule assembly. Physiol Rev. 1995;75(4):835–64.

31. Stein GS, Lian JB, Stein JL, Van Wijnen AJ, Montecino M. Transcriptional control of osteoblast growth and differentiation. Physiol Rev. 1996;76(2):593–629.

32. Zhao F, van Rietbergen B, Ito K, Hofmann S. Fluid flow-induced cell stimulation in bone tissue engineering changes due to interstitial tissue formation in vitro. Int J Numer Method Biomed Eng. 2020;36(6):e3342.

33. Bancroft GN, Sikavitsas VI, Mikos AG. Design of a flow perfusion bioreactor system for bone tissue-engineering applications. Tissue Eng. 2003;9(3):549–54.

34. Martin I, Wendt D, Heberer M. The role of bioreactors in tissue engineering. Trends Biotechnol. 2004;22(2):80–6.

35. McCoy RJ, O’Brien FJ. Influence of shear stress in perfusion bioreactor cultures for the development of three-dimensional bone tissue constructs: a review. Tissue Eng Part B Rev. 2010;16(6):587–601.

36. Nguyen BN, Ko H, Moriarty RA, Etheridge JM, Fisher JP. Dynamic Bioreactor Culture of High Volume Engineered Bone Tissue. Tissue Eng Part A. 2016;22(3-4):263–71.

